# Rare variants in the genetic background modulate the expressivity of neurodevelopmental disorders

**DOI:** 10.1101/257758

**Authors:** Lucilla Pizzo, Matthew Jensen, Andrew Polyak, Jill A. Rosenfeld, Katrin Mannik, Arjun Krishnan, Elizabeth McCready, Olivier Pichon, Cedric Le Caignec, Anke Van Dijck, Kate Pope, Els Voorhoeve, Jieun Yoon, Paweł Stankiewicz, Sau Wai Cheung, Damian Pazuchanics, Emily Huber, Vijay Kumar, Rachel Kember, Francesca Mari, Aurora Curró, Lucia Castiglia, Ornella Galesi, Emanuela Avola, Teresa Mattina, Marco Fichera, Luana Mandarà, Marie Vincent, Mathilde Nizon, Sandra Mercier, Claire Bénéteau, Sophie Blesson, Dominique Martin-Coignard, Anne-Laure Mosca-Boidron, Jean-Hubert Caberg, Maja Bucan, Susan Zeesman, Małgorzata J.M. Nowaczyk, Mathilde Lefebvre, Laurence Faivre, Patrick Callier, Cindy Skinner, Boris Keren, Charles Perrine, Paolo Prontera, Nathalie Marle, Alessandra Renieri, Alexandre Reymond, R Frank Kooy, Bertrand Isidor, Charles Schwartz, Corrado Romano, Erik Sistermans, David J. Amor, Joris Andrieux, Santhosh Girirajan

## Abstract

**Purpose:** To assess the contribution of rare variants in the genetic background towards variability of neurodevelopmental phenotypes in individuals with rare copy-number variants (CNVs) and gene-disruptive mutations.

**Methods:** We analyzed quantitative clinical information, exome-sequencing, and microarray data from 757 probands and 233 parents and siblings who carry disease-associated mutations.

**Results:** The number of rare secondary mutations in functionally intolerant genes (second-hits) correlated with the expressivity of neurodevelopmental phenotypes in probands with 16p12.1 deletion (n=23, p=0.004) and in probands with autism carrying gene-disruptive mutations (n=184, p=0.03) compared to their carrier family members. Probands with 16p12.1 deletion and a strong family history presented more severe clinical features (p=0.04) and higher burden of second-hits compared to those with mild/no family history (p=0.001). The number of secondary variants also correlated with the severity of cognitive impairment in probands carrying pathogenic rare CNVs (n=53) or *de novo* mutations in disease genes (n=290), and negatively correlated with head size among 80 probands with 16p11.2 deletion. These second-hits involved known disease-associated genes such as *SETD5, AUTS2*, and *NRXN1*, and were enriched for genes affecting cellular and developmental processes.

**Conclusion:** Accurate genetic diagnosis of complex disorders will require complete evaluation of the genetic background even after a candidate gene mutation is identified.

## Introduction

Significant advances in high-throughput genomic sequencing technologies have helped to identify hundreds of genes as risk factors for neurodevelopmental and neuropsychiatric disorders, including autism, intellectual disability, schizophrenia, and epilepsy. For example, in 2002, only 2-3% of autism cases were explained by genetic factors, whereas current studies suggest that rare disruptive mutations, including copy-number variants (CNVs) and single-nucleotide variants (SNVs), account for 10-30% of autism cases^1^. Despite initial claims of association with a specific disorder or syndrome, several of these pathogenic variants show incomplete penetrance and variable expressivity^2–4^. For example, the 16p11.2 BP4-BP5 deletion (OMIM #611913) was first described in children with autism, but further studies on other clinical and population cohorts demonstrated that this deletion is also associated with intellectual disability and developmental delay (ID/DD), obesity, epilepsy, cardiac disease, and scoliosis, and only about 24% of cases manifest an autism phenotype^5–9^. Phenotypic variability is not restricted to multi-genic CNVs but has also been reported for single genes with disruptive mutations, including *DISC1, PTEN, SCN1A, CHD2, NRXN1, FOXP2*, and *GRIN2B*3. While some of these effects could be due to allelic heterogeneity, phenotypic variability among carriers of the same molecular lesion suggests a strong role for variants in the genetic background^10,11^. For example, in a large family described by St. Clair and colleagues, carriers of a balanced translocation disrupting *DISC1* manifested a wide range of neuropsychiatric features, including schizophrenia, bipolar disorder and depression^12^.

This phenomenon was exemplified by our delineation of a 520-kbp deletion on chromosome 16p12.1 (OMIM #136570) that is associated with developmental delay and extensive phenotypic variability^13^. Interestingly, in most cases, this deletion was inherited from a parent who also manifested mild neuropsychiatric features, and the severely affected children were more likely to carry another large (>500 kbp) rare CNV. We hypothesized that while each pathogenic primary mutation sensitizes the genome to varying extents, additional secondary variants in the genetic background modulate the ultimate clinical manifestation.

While previous studies have only assessed the role of large CNV second hits towards variability of disease-associated CNVs^2^, the combined effect of SNVs, insertion/deletions and CNVs in the genetic background towards variability of specific phenotypes among individuals carrying mutations (CNVs or SNVs) associated with neurodevelopmental disease has not been assessed. In this study, we evaluated 757 probands and 233 family members carrying disease-associated primary mutations (17 rare CNVs or disruptive mutations in 301 genes). A comparison of the genetic background between probands and parents or siblings showed that in the presence of the same primary mutation, variability and severity of neurodevelopmental disease correlates with the number of rare pathogenic variants, suggesting a global role of the genetic background towards phenotypic heterogeneity.

## Materials and methods

### Cohorts analyzed

We analyzed clinical and genetic data in individuals carrying a disease-associated “primary variant” or “first-hit”, defined as follows: (a) rare CNVs previously associated with neurodevelopmental and neuropsychiatric disorders^2^, (b) previously reported *de novo* mutations in candidate genes^14,15^, and (c) likely damaging variants in recurrent genes associated with neurodevelopmental and neuropsychiatric disorders. Five subgroups of individuals carrying a disease-associated primary variant were analyzed (Figure 1): (a) 26 probands, 23 carrier parents and available family members carrying 16p12.1 deletion, (b) 53 autism probands from the Simons Simplex Collection (SSC) cohort who carry rare CNVs associated with syndromic and variably expressive genomic disorders^2^, (c) 84 probands and available family members with 16p11.2 BP4-BP5 deletion from the Simons Variation in Individuals Project (SVIP) cohort, (d) 295 autism probands from the SSC cohort reported to carry severe *de novo* loss-of-function mutations in neurodevelopmental genes^14,15^, (e) 184 probands and matched unaffected siblings from the SSC cohort who inherited the same rare (≤0.1% frequency) loss-of-function or likely pathogenic missense primary mutations (CADD ≥25) in genes recurrently disrupted in neurodevelopmental disorders^16^.

**Figure 1.**
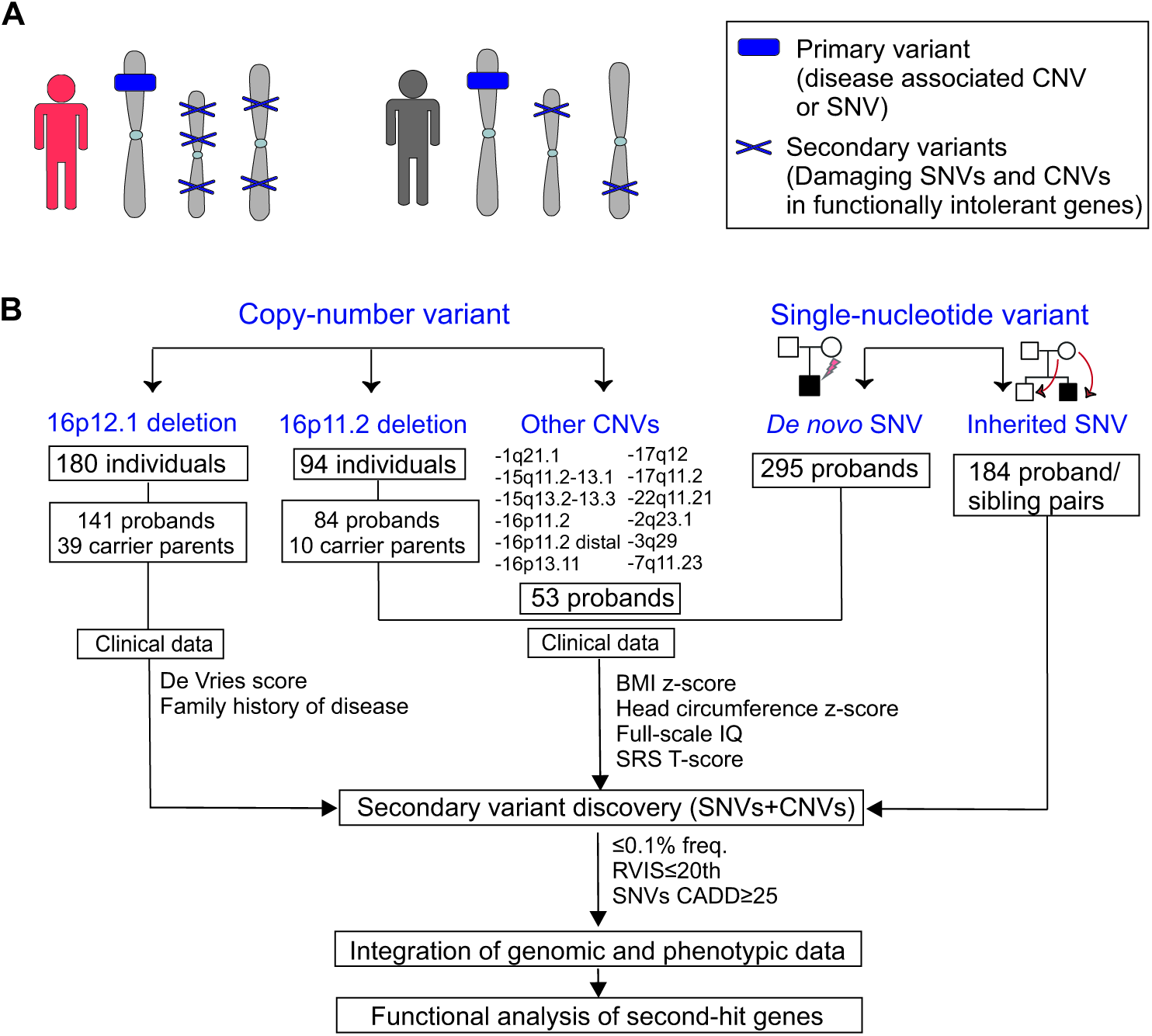
Strategy for understanding the role of the genetic background in phenotypic variability of neurodevelopmental disease. (**A)** Schematic of primary and secondary variants used in this study. Disease-associated mutations common among different individuals were considered as “primary variants” (blue box), and rare likely pathogenic SNVs or CNVs affecting functionally intolerant genes were defined as “secondary variants” (blue Xs). Individuals with a high burden of secondary variants (in red) exhibit a more severe clinical manifestation compared to those carrying the same primary variant but with a lower number of second-hits (in gray). (**B)** Combined clinical and genomic analysis of 757 probands and 233 family members carrying primary disease-associated variants (16p12.1 deletion, 16p11.2 deletion, 16 rare CNVs, *de novo* mutations in autism simplex cases, and inherited mutations in disease-associated genes) was performed to understand the role of rare (<0.1%) likely pathogenic secondary variants (SNVs with CADD≥25 and CNVs) in functionally intolerant genes (RVIS≤20th percentile) towards thevariable manifestation of neurodevelopmental disease.

### Patient recruitment and clinical data ascertainment

We obtained clinical and/or genomic data from 141 children carrying the 16p12.1 deletion, as well as 39 deletion carrier and 30 non-carrier parents. Probands and parents recruited through direct contact provided consent according to the protocol reviewed and approved by The Pennsylvania State University Institutional Review Board. When individuals were not contacted directly, de-identified phenotypic (case histories and clinical information) and genomic data were used; as such, these cases were exempt from IRB review and conformed to the Helsinki Declaration.

We extracted clinical information from medical records or clinical questionnaires completed by different physicians from 180 carrier individuals and available family members (Supplementary Method 1.1). We used a modified de Vries scoring system for quantifying the number and severity of phenotypic abnormalities in affected children, which allows for a uniform assessment of developmental phenotypes from clinical records (Table S1)^17^. Family history information was used to bin families with the deletion into strong and mild or negative family history categories based on the severity of neurodevelopmental or psychiatric features (Figure S1, Supplementary Method 1.1). Genomic and clinical data for SSC and SVIP cohort families were obtained from the Simons Foundation Autism Research Initiative (SFARI) following appropriate approvals (see Supplementary Method 1.1).

### Secondary variant burden analysis

To identify all coding variants modulating the presentation of the 16p12.1 deletion, we performed exome sequencing and SNP arrays on 105 individuals from 26 families as previously reported (Supplementary Method 1.2). Variant calls (SNVs and CNVs) from 716 individuals in the SSC were obtained from exome and SNP microarray studies^14,15,18^, and variant call files (VCF) and SNP array data from 84 families with 16p11.2 BP4-BP5 deletion were obtained from the Simons Foundation. Single-nucleotide variants and CNVs were filtered following the same procedures as for the 16p12.1 deletion cohort (Supplementary Method 1.2). We defined “secondary variant” or “second-hit” as an additional rare likely pathogenic mutation (<0.1% frequency CNV or SNV with CADD≥25) affecting a functionally intolerant gene (RVIS≤20^th^ percentile) in an individual who already carries a disease-associated primary mutation (Figure 1A, Supplementary Method 1.2). The function of second-hit genes was analyzed using Ingenuity Pathway Analysis (IPA, Qiagen Bioinformatics), expression data derived from the GTEx consortium^19^, and Gene ontology (GO) enrichment analysis^20^ (Supplementary Method 1.3).

### Statistical analyses

Proband-parent or proband-sibling secondary variant burdens and clinical severity scores were compared using Wilcoxon signed-rank test. Genetic burden and clinical severity scores between different categories of probands with related or shared primary variants were compared using non-parametric one-tailed Mann-Whitney tests, due to the hypothesis-driven nature of the comparison. The Kolmogorov-Smirnov test was used to assess normality distribution of the burden of secondary variants, FSIQ, SRS T-scores, BMI z-scores and head circumference (HC) z-scores. Correlation between number of secondary variants and quantitative phenotypes was assessed using Pearson’s correlation for normally distributed datasets, or Spearman’s correlation for datasets that were not normally distributed. Statistics were calculated using Minitab or R (v.3.4.2) software. Boxplots display the distribution of data from minimum to maximum.

## Results

### Secondary variants and disease expressivity in 16p12.1 deletion probands

We assessed how rare variants in the genetic background can modulate phenotypes in concert with a first-hit by evaluating 757 affected probands and 233 family members carrying disease-associated primary variants (rare CNVs or disruptive gene mutations) (Figure 1, see Methods). Using the 16p12.1 deletion as a paradigm for studying the genetic basis of variable expression of disease traits, we analyzed 180 individuals with the deletion and their non-carrier family members (Figure 1B). The 16p12.1 deletion was inherited in 92.4% of cases, with a significant maternal bias (57.6% maternal (n=53) vs 34.8% paternal (n=32), one-tailed binomial test p=0.01) (Table S2). In accordance with the female protective model described for neurodevelopmental disorders^2,21,22^, we observed a significant gender bias among probands with the 16p12.1 deletion (67.9% males versus 32.1% females, one-tailed binomial test p<0.0001). Detailed clinical analysis of 141 affected children with 16p12.1 deletion showed a wide heterogeneity of phenotypes, with a high prevalence of neurodevelopmental, craniofacial and musculoskeletal features (>50%), and variable involvement of other organs and systems (Figure 2A, Table S3). In contrast, 32 of 39 (82%) (61.5% females, 38.5% males) carrier parents showed mild cognitive, behavioral and/or psychiatric features (Table S4), consistent with previous reports of cognitive impairment and increased risk for schizophrenia in carriers of the 16p12.1 deletion^23,24^.

**Figure 2.**
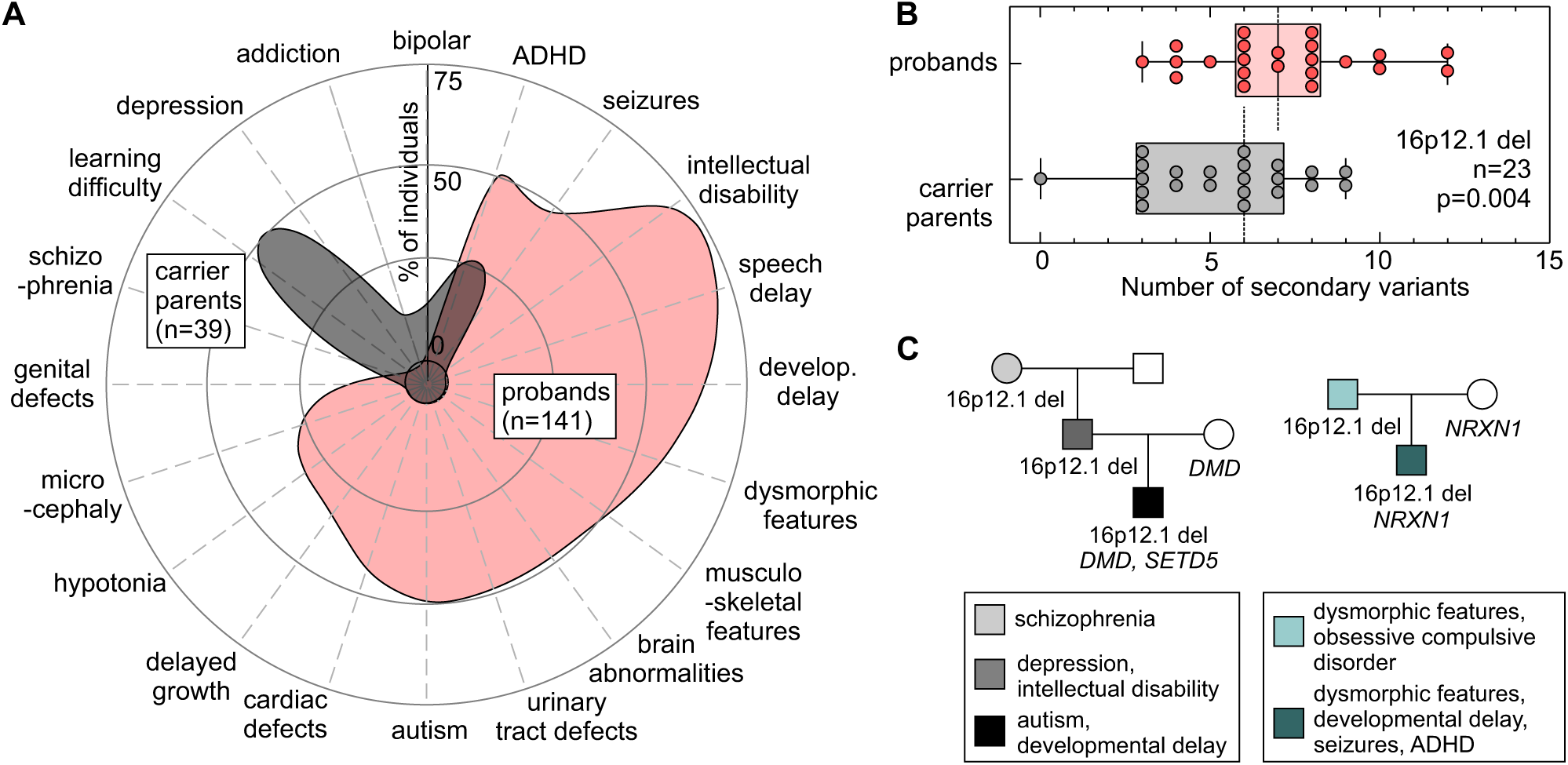
Rare pathogenic mutations in functionally intolerant genes contribute to the phenotypic heterogeneity in 16p12.1 deletion. (**A)** Phenotypic spectrum of 16p12.1 deletion in probands (n=141, red) and carrier parents (n=39, grey). Probands exhibit a spectrum of severe developmental features compared to the mild cognitive and psychiatric features observed in carrier parents. Features represented were observed in ≥5% of probands or carrier parents. (**B)** Analysis of rare (≤0.1%) likely pathogenic secondary variants (SNVs with CADD≥25) in genes intolerant to functional variation (RVIS≤20th percentile) in proband-carrier parent pairs shows that probands present a higher second-hit mutation burden compared to their carrier parents (n=23, Wilcoxon signed-rank test, p=0.004). (**C)** Example of families with inherited 16p12.1 deletion. Family 1 shows three generations carrying 16p12.1 deletion and multiple neurodevelopmental and psychiatric features, with the proband carrying a *de novo* LoF mutation in *SETD5* (p.Asp542Thrfs*3) and a stopgain mutation in *DMD* gene (p.Trp3X) inherited from the mother without the 16p12.1 deletion (non-carrier). Family 2 shows a proband with multiple congenital and neurodevelopmental features carrying 16p12.1 deletion and 2p16.3 deletion (encompassing *NRXN1*), the latter inherited from the non-carrier mother.

To identify variants within protein-coding regions that contribute to a more severe manifestation of the deletion in the affected children compared to their carrier parents, we performed exome sequencing and high-resolution SNP arrays in 26 families (n=105) with 16p12.1 deletion (23 inherited and 3 *de novo* cases, Table S5). We first evaluated whether the deletion could unmask recessive alleles, and found no rare pathogenic mutations within the seven 16p12.1 genes on the non-deleted chromosome (Table S6). We next performed a case-by-case analysis of families for second hits elsewhere in the genome by focusing on rare CNVs (<0.1%, ≥50 kbp), *de novo* or rare (ExAC frequency <0.1%) loss-of-function (LoF) mutations, and rare likely pathogenic missense variants (Phred-like CADD ≥25) in disease-associated genes (see Methods, Tables S7-S9). For example, in one proband we identified two disease-associated mutations, including a *de novo* LoF mutation in the intellectual disability-associated gene *SETD5* (OMIM #615761, c.1623_1624insAC, p.Asp542Thrfs*3) and a LoF mutation in *DMD* (OMIM #310200, c.9G>A, p.Trp3X) transmitted from the 16p12.1-deletion carrier mother (Figure 2C). Similarly, a rare deletion at 2p16.3 encompassing *NRXN1* (OMIM #614332), inherited from the non-carrier mother, was identified in another proband (Figure 2C).

While private disease-associated secondary mutations may explain the variable and severe features in the affected children on a case-by-case basis, we lacked the statistical power to implicate individual genes or variants that modulate specific 16p12.1 deletion phenotypes. Therefore, to globally assess the genome-wide contribution of rare pathogenic mutations affecting functionally relevant genes, we performed an integrative analysis and quantified rare (frequency ≤ 0.1%), likely pathogenic variants (CNVs or SNVs with CADD≥25) within genes intolerant to functional variation. The Residual Variation Intolerance Score (RVIS) has been shown to be a good predictor of intolerance of a gene to pathogenic mutations, and has been widely used by multiple studies for the recapitulation of known and the discovery of novel disease-associated genes^15,25^. For example, Krumm *et al*. showed that autism genes have an average RVIS of 26^th^ percentile, and using classifications from the human disease network, Petrovski *et al*. showed that genes involved in developmental, skeletal, cardiac, neurological and muscular disorders show average scores in the 20^th^-30^th^ percentile^15,26^. Therefore, to focus our assessment of secondary mutations to a subset of genes relevant to disease pathogenicity, we quantified rare likely pathogenic variants in genes with RVIS≤20^th^ percentile, hereafter referred to as the “burden of secondary variants” or “second hits”. Intra-familial comparison of second-hit burden showed that probands have an excess of secondary variants compared to their carrier parents (Wilcoxon signed-rank test, p=0.004, Figures 2B and S2A-D), with no change in the number of synonymous mutations in all genes (p=0.29) or in RVIS≤20^th^ genes (p=0.36, Figures S2E-F). Further, functional analysis of second-hit genes showed that probands presented an excess of second-hit genes that were preferentially expressed in the human brain (Wilcoxon signed-rank test, p=0.04, Figure S3) and enriched for developmental pathways (Table S10) compared to carrier parents.

The severity and variability of neurodevelopmental features is contingent upon family history of neuropsychiatric disease^22^. In fact, the cognitive and social outcomes in probands with *de novo* 16p11.2 BP4-BP5 deletion or 22q11.2 deletion have been reported to positively correlate with the cognitive and social skills of their parents^27,28^. However, the genetic basis of such background effects has not been specifically studied. We assessed the role of second-hit burden in the family-specific background effects and in the observed inter-familial variability of clinical features in probands with 16p12.1 deletion. We found that probands with a strong family history of neurodevelopmental and psychiatric disease presented a more severe and heterogeneous clinical presentation (Mann Whitney one-tailed, p=0.04) and a higher burden of secondary variants (p=0.001) than those with mild or negative family history (Figures 3A-C and S4A-C). Interestingly, probands with a strong family history also showed a higher difference in burden compared to their carrier parents than probands with a mild family history (p=0.003, Figure 3D). While we did not observe a difference in burden between carrier parents based on family history (p=0.68, Figure S4B), we found that non-carrier parents with a strong family history presented a significantly higher burden compared to those with a mild family history (p=0.01, Figure 3E). Therefore, in families with a strong history of neurodevelopmental and psychiatric disease, a higher number of rare and likely pathogenic secondary variants are more likely to be transmitted to the proband from the non-carrier parent, potentially contributing to a more severe manifestation of the disorder. These results suggest a potential role for rare pathogenic variants in the genetic background in modulating intra- and inter-familial clinical variability observed in families with the 16p12.1 deletion.

**Figure 3:**
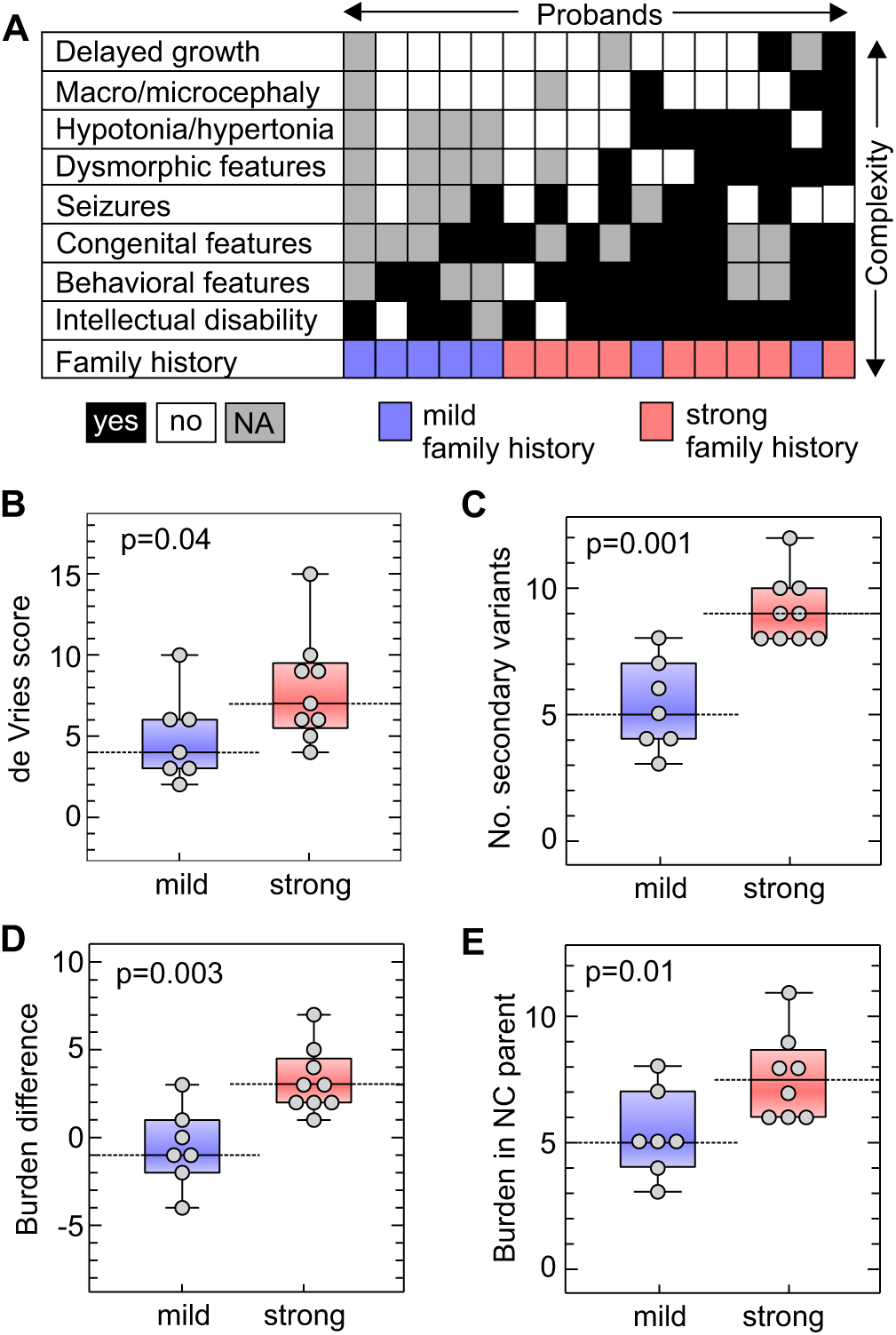
Strong family history of neurodevelopmental and psychiatric disease is associated with excess of secondary variants and severe phenotypic outcome in 16p12.1 deletion probands. (**A)** Diagram showing phenotypic heterogeneity in 16 probands with 16p12.1 deletion (black= phenotype present, white=absent, grey=not assessed) and their family history of neurodevelopmental and psychiatric disease (red=strong, blue=mild/negative). Probands with strong family history (n=9) have (**B)** a more heterogeneous clinical manifestation (higher de Vries scores, one-tailed, Mann-Whitney, p=0.04) and (**C)** a higher burden of secondary variants (one-tailed Mann-Whitney p=0.001) than those with mild or negative family history (n=7). (**D)** Probands with a strong family history exhibit a greater difference in second-hit burden compared to carrier parents (p=0.003). (**E)** Non-carrier parents from families with strong family history present a higher burden compared to those with mild/negative family history (one-tailed Mann-Whitney, p=0.01).

### Secondary variant burden correlates with quantitative phenotypes among individuals with 16p11.2 deletion and other rare pathogenic CNVs

We next assessed whether the second-hit burden modulates quantitative phenotypes in carriers of other CNVs associated with neurodevelopmental phenotypes (Figure 1B). In probands with disease-associated rare CNVs (n=53, Table S11) from the Simons Simplex Cohort (SSC), we observed a modest but significant negative correlation (Pearson correlation, R=−0.36, p=0.004) between the number of secondary variants and full-scale IQ scores (FSIQ). This result held true when we separately analyzed individuals carrying 16p11.2 BP4-BP5 deletion (R=−0.68, p=0.04), but did not show statistical significance for 16p11.2 BP4-BP5 duplication (R =−0.34, p=0.17), 1q21.1 duplication (R =−0.36, p=0.32), or 7q11.23 duplication (R =−0.74, p=0.17), potentially due to a low sample size (Figures 4A and S5). Interestingly, probands with disease-associated CNVs and intellectual disability (FSIQ<70) showed a significant increase in the number of secondary variants compared to those without intellectual disability (FSIQ≥70, one-tailed Mann-Whitney, p=0.02, Figure S6).

**Figure 4.**
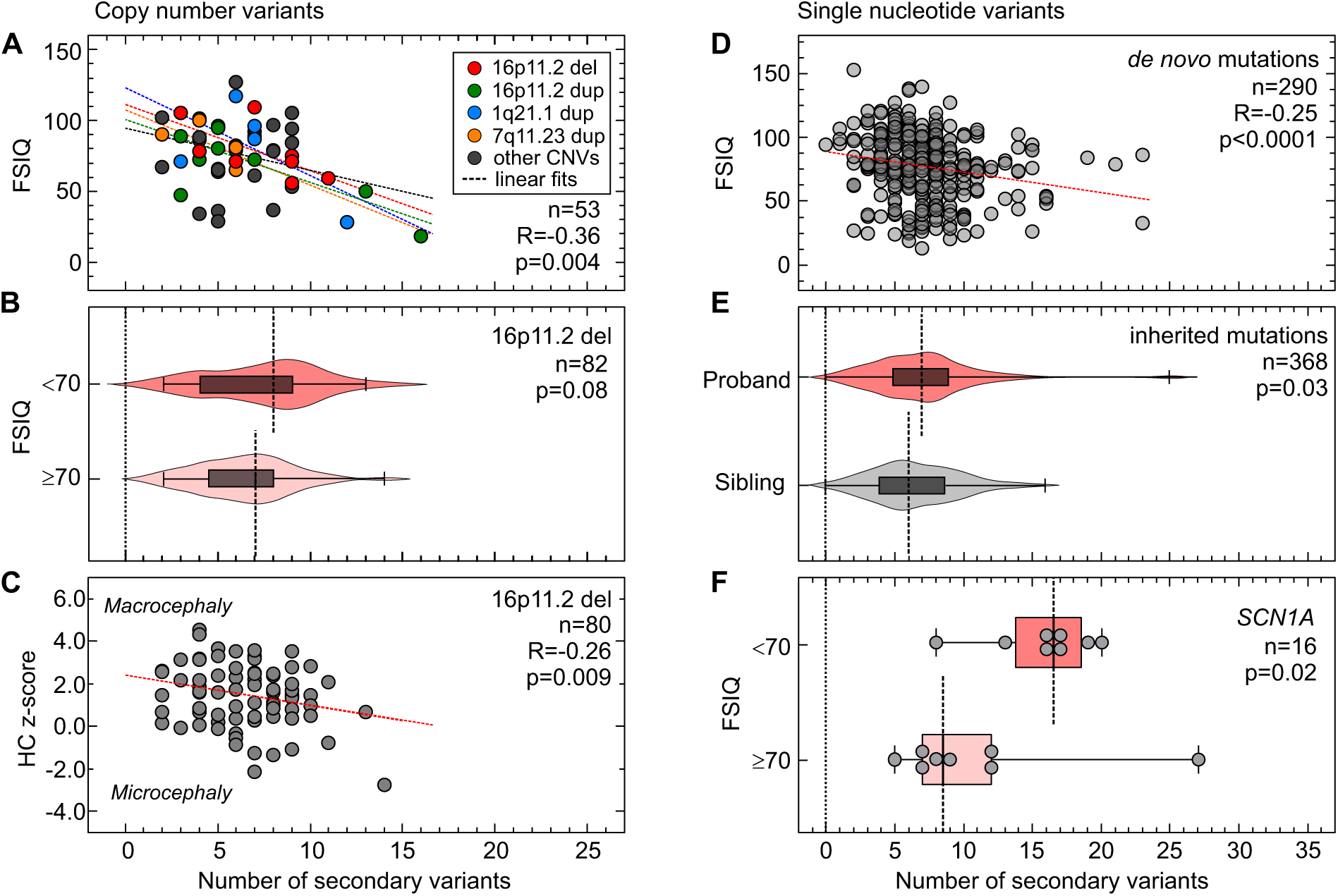
Burden of secondary variants modulates quantitative phenotypes among probands with a first-hit CNV or SNV associated with neurodevelopmental disease. (**A)** Negative correlation between the number of secondary variants and full-scale IQ (FSIQ) scores in individuals (n=53) carrying 16 CNVs associated with neurodevelopmental disease (Pearson correlation, R=−0.36, p=0.004). Probands with 16p11.2 deletion (red), 16p11.2 duplication (green), 1q21.1 duplication (blue) and 7q11.23 duplication (yellow) are highlighted, while grey circles represent probands with other rare CNVs. (**B)** Higher burden of secondary variants among probands with 16p11.2 deletion and FSIQ<70 (n=17) compared to probands with FSIQ≥70 (n=65, one-tailed, Mann-Whitney, p=0.08). (**C)** Negative correlation between the number of secondary variants and head circumference z-scores (age≥12 months, n=80, Pearson correlation R=−0.26, p=0.009) in probands with 16p11.2 deletion. (**D)** Autism probands with *de novo* disruptive mutations and available FSIQ scores (n=290) show a moderate negative correlation (Spearman correlation coefficient, R=−0.25, p<0.0001) between the number of secondary variants and FSIQ scores. (**E)** Probands present an excess of secondary variants compared to their unaffected siblings (n=184 pairs) carrying the same inherited pathogenic mutations (loss-of function or missense CADD≥25) in genes recurrently disrupted in neurodevelopmental disease (Wilcoxon signed-rank test, p=0.03). (**F)** Enrichment of secondary variants among individuals with damaging mutations in *SCN1A* (loss-of-function or missense CADD≥25) and intellectual disability (one-tailed Mann-Whitney, p=0.02) compared to those without intellectual disability.

We further expanded our analysis by evaluating a larger set of 84 families with 16p11.2 BP4-BP5 deletion from the Simons Variation in Individuals Project (SVIP). We observed a higher median number of secondary variants in probands carrying the 16p11.2 deletion that had intellectual disability (FSIQ<70, median=8) compared to those with no intellectual disability (FSIQ≥70, median=7 one-tailed Mann-Whitney, p=0.08, Figure 4B), without a difference in the number of synonymous mutations between the two subgroups (median of 9,957 synonymous changes for FSIQ<70 group versus 10,052 for the FSIQ≥70 group, two-tailed Mann-Whitney, p=0.51, Figure S7A). Notably, we observed only a mild negative correlation between second-hit burden and FSIQ, which did not attain statistical significance (Pearson correlation, R=−0.16, p=0.08, Figure S7B). We hypothesized that this marginal significance compared to 16p11.2 deletion probands from the SSC cohort (Figures 4A and S5B) could be due to differences in clinical ascertainment, as the SVIP cohort was selected for individuals carrying a 16p11.2 deletion who manifested a more heterogeneous set of phenotypes, while individuals from the SSC cohort were specifically ascertained for idiopathic autism^29^. These differences in ascertainment were evident by different distributions of quantitative phenotypes, including BMI, FSIQ, and SRS T-scores, in both populations (see Figure S8).

After adjusting for age to allow for full manifestation of the head phenotype, we identified a negative correlation between the number of secondary variants in SVIP probands with 16p11.2 deletion and their head circumference (HC) z-scores (age≥12 months, n=80, Pearson’s R=−0.26, p=0.009, Figure 4C)^8,30^. The observation that HC z-scores decline steadily (from >2 to <-2 scores) as second hits accumulate confirms that the deletion primarily leads to macrocephaly phenotypes, and suggests that the second-hits could explain the incomplete penetrance of this phenotype among carriers of the deletion^30^. We note that the second-hit burden did not correlate with Social Responsiveness Scale (SRS) T-scores or Body-Mass Index (BMI) z-scores, measures for autism and obesity, among probands with rare CNVs from the SSC cohort (Figures S9A-B) or those with 16p11.2 deletion from the SVIP cohort (Figures S9C-D), suggesting other mechanisms for the variability of these phenotypes.

### Secondary variants modulate disease manifestation among individuals with disruptive mutations in disease-associated genes

We next analyzed 295 simplex cases from the SSC cohort with previously identified *de novo* gene-disruptive mutations within 271 genes^14,15^, and observed a moderately negative correlation between secondary variant burden and FSIQ scores (Spearman’s correlation, R=−0.25, p<0.0001, Figure 4D). Within this cohort, individuals with intellectual disability (FSIQ<70, n=93) presented an enrichment of second hits compared to those without intellectual disability (FSIQ≥70, n=197) (one-tailed Mann-Whitney, p=0.001, Figure S10A). We did not observe a role for secondary variant burden in modulating BMI z-scores (Spearman’s R=−0.038, p=0.27, Figure S10B), although we did find a mild but significant positive correlation with SRS T-scores (Spearman’s R=0.12, p=0.02, Figure S10C). Moreover, when probands were separated by gender, we observed a higher burden of secondary variants in females compared to males (one-tailed Mann-Whitney, p=0.02, Figure S11). This supports the hypothesis that females require a higher contribution from the genetic background to reach the genetic threshold for pathogenesis of neurodevelopmental disease than males^22^.

While there is a consensus on the pathogenic role of *de novo* gene-disruptive mutations in simplex families, the interpretation of inherited disruptive variants within the same genes is challenging. To understand the role of the genetic background in the penetrance of inherited disruptive mutations in disease-associated genes, we analyzed 184 pairs of probands and unaffected siblings who inherited the same pathogenic mutations in genes recurrently disrupted in neurodevelopmental disorders (Table S12). We found a greater enrichment of second hits in probands compared to their unaffected siblings (Wilcoxon signed-rank test p=0.03, Figure 4E), suggesting that second hits likely contribute to increased penetrance of neurodevelopmental phenotypes in children with inherited pathogenic single-gene mutations. When we analyzed probands carrying disruptive mutations in specific neurodevelopmental genes, we found that the severity of cognitive deficits in individuals with damaging mutations in *SCN1A* was concordant with an excess of secondary variants (probands with FSIQ<70, n=8, median=16.5, versus those with FSIQ≥70, n=8, median=8.5, one-tailed Mann-Whitney, p=0.02) (Figure 4F). This observation could also explain the diversity of other phenotypes co-occurring with the disruption of the epilepsy-associated *SCN1A* gene, such as motor delay and autism^31^.

### Secondary variants involve disease associated genes and affect core cellular and developmental processes

To understand how secondary variants could modulate phenotypes among probands with pathogenic first-hit variants, we explored the functionality of second-hit genes identified in all probands analyzed in our study. Overall, we identified 3,197 rare likely-pathogenic mutations encompassing a diverse set of 1,615 functionally intolerant genes. Of these, 40.9% (660/1,615) were found to be extremely intolerant to loss-of-function mutations (pLI metric ≥0.9). These genes were also enriched for postsynaptic density genes, genes encoding FMRP protein targets, chromatin-associated genes, genes embryonically expressed, and essential genes, compared to the whole genome (52% vs. 26%, Chi-squared test, p<0.0001)^32,33^. Interestingly, 44 of these second-hit genes (such as *CNTNAP2, MBD5, SCN1A, CHD8* and *AUTS2*) have been recurrently associated with neurodevelopmental disorders^16^ (Figure S12), 58 genes have been previously identified as a causative gene in simplex autism cases^14,15^ (Table S13), and 50 genes have been associated with skeletal, muscular, cardiovascular or renal disorders, as classified in the human disease network from the OMIM database (Table S14)^34^. We further assessed the location of second-hit mutations within a subset of genes recurrently associated with disease, including *RIMS1, DIP2A, KDM5B* and *ACOX2*, and found no specificity for the location of the secondary variants within the protein sequences compared to previously reported *de novo* disruptive mutations within these genes (Figure 5A). In fact, in some cases, we observed stopgain secondary variants that were more premature in the protein sequence than previously reported disruptive mutations in these genes, suggesting that the second-hit can potentially exert an effect as severe as the first-hit, if these are true loss-of-function variants. The allelic diversity of second hits within these genes suggests that further functional analysis should be performed in order to understand their specific effects on modulating developmental phenotypes.

**Figure 5.**
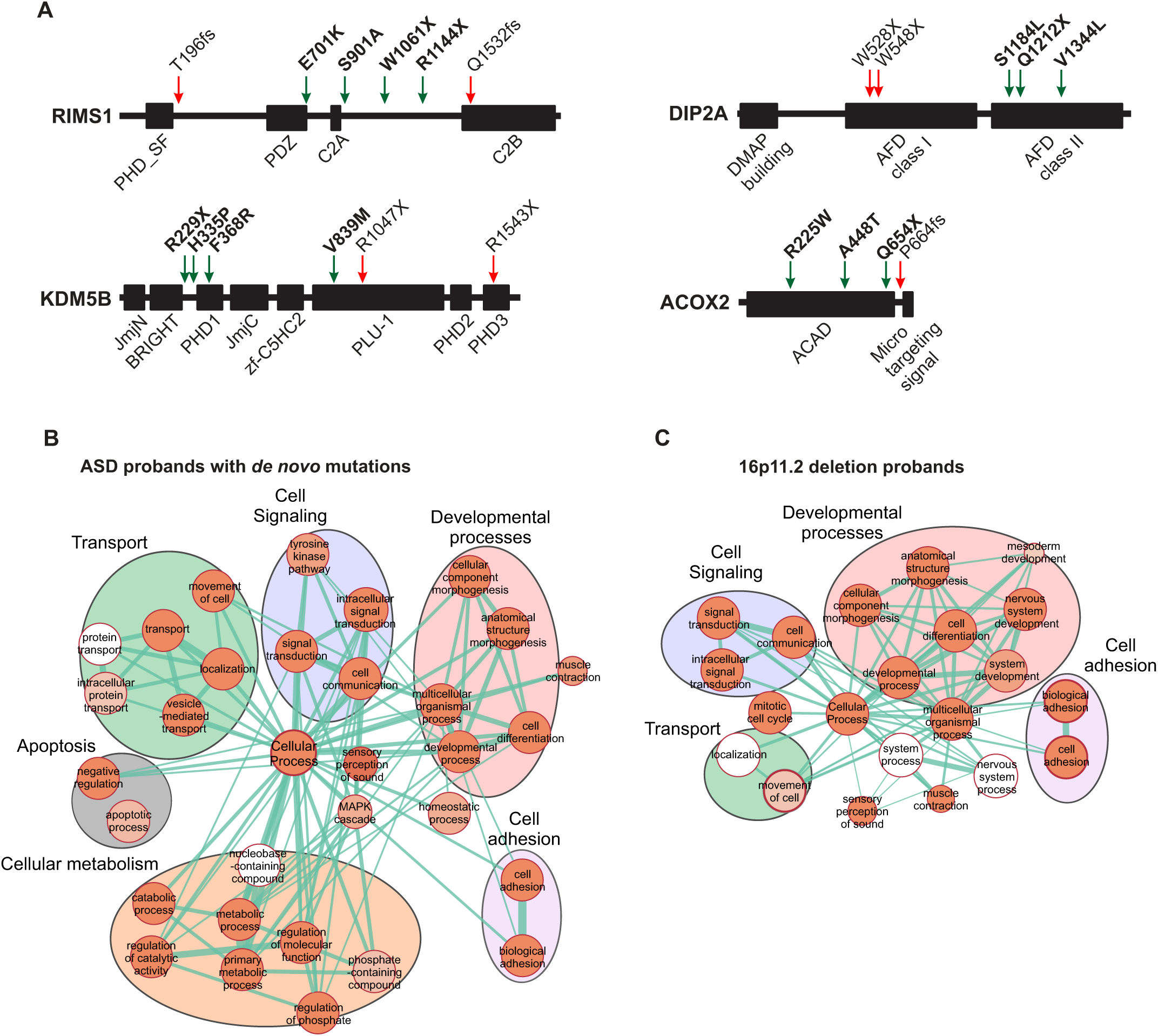
Secondary variants affect core biological processes and affect disease-associated genes. (**A)** Examples of non-specificity in the location of secondary variants in protein domains compared to first-hits. Location of variants in the protein sequences of *RIMS1, DIP2A, KDM5B* and *ACOX2*, recurrent genes with secondary variants (green arrows) in probands with first-hit pathogenic CNVs or *de novo* disruptive mutations and previously reported to carry *de novo* disrupting mutations in simplex autism cases (red arrows). Genes with secondary variants found in (**B)** autism probands carrying *de novo* disruptive mutations (SSC) and (**C)** probands with the 16p11.2 deletion (SVIP) are enriched in core biological processes (FDR<0.05 with Bonferroni correction). Recurrent clusters of enriched gene ontology terms for “developmental processes”, “cell signaling”, “cell adhesion” and “transport” functions are present among second hits found in each cohort. The size of each circle represents the number of genes annotated for each GO term; red shading of each circle represents the FDR for enrichment of each GO term among second-hit genes in each cohort, with darker shades indicating a lower FDR. Line thickness represents the number of shared annotated genes between pairs of GO terms.

To further characterize the function of second-hit genes, we performed Gene Ontology enrichment analysis of second-hit genes in probands with *de novo* mutations from the SSC and 16p11.2 deletion probands from the SVIP cohort. We found that genes carrying secondary variants in probands from both cohorts were enriched for core processes, including cell signaling and adhesion and developmental processes (Figures 5B-C, Table S15-S16). Although some of these genes have been individually associated with a disease phenotype, further functional analyses are required to understand potential interactions between genes affected by primary and secondary variants, their cumulative burden, and ultimately their potential contribution towards phenotypic variability.

## Discussion

Our results support an oligogenic model for neurodevelopmental disorders, where the primary variant provides a sensitized genomic background for disease and secondary variants within functionally intolerant genes modify the phenotypic trajectory of the disorder. We propose that a higher burden of secondary variants increases the likelihood of involving a disease-associated modifier gene within disease-related pathways, as well as allows for a higher number of oligogenic combinations potentially modulating the phenotype associated with the first-hit. Some primary variants that are more tolerant to changes in the genetic background, such as the 16p12.1 deletion, are transmitted through generations and only surpass the threshold for severe disease with the accumulation of several rare pathogenic mutations. Other primary variants which are often *de novo*, such as the 16p11.2 deletion, push the genetic background closer to the threshold for severe manifestation and therefore require a lesser contribution from secondary variants. Similarly, highly penetrant syndromic CNVs such as Smith-Magenis syndrome and Sotos syndrome, which are mostly *de novo* and encompass genes more intolerant to functional variation compared to variably expressive CNVs (p=0.03, one-tailed Mann-Whitney, Table S17, Figure S13), would push the genetic liability beyond the threshold for severe disease^2,13^. While additional mutations may not be necessary for complete penetrance of these disorders, these variants can modify specific phenotype traits when present. For example, deleterious mutations in histone modifier genes have been reported to contribute to the conotruncal heart manifestation of 22q11.2 deletion syndrome^35^. This model would also apply for single-gene disorders, where second hits potentially explain discordant clinical features reported among affected carriers of the same molecular alteration, as described for Rett syndrome and individuals with disruptive mutations in the intellectual disability gene *PACS1*^36,37^.

Our observations that probands with a strong family history exhibit severe clinical manifestation and a higher second-hit burden also provide insights into the role of rare variants in the genetic background towards the reported correlation between parental profiles and clinical outcome in probands carrying rare CNVs^27,28^. Moreover, the observed higher burden of secondary variants in the non-carrier parents in families with strong family history suggests assortative mating, and transmission of these hits to the proband potentially explains the increased clinical severity. Similarly, in 16p11.2 deletion, we observed that children who inherited the CNV presented lower FSIQ scores (n=8, median FSIQ=75) than probands with a *de novo* deletion (n=57, median FSIQ=85, one-tailed Mann-Whitney, p=0.006, Figure S14A-B), in agreement with previous reports^8^. This was also consistent with a non-significant excess of the second-hit burden among probands with an inherited 16p11.2 deletion (one-tailed Mann-Whitney, p=0.06, Figure S14B) compared to *de novo* probands. These results highlight the importance of eliciting family history of psychiatric and neurodevelopmental disease for more accurate diagnostic assessment of the affected children.

Overall, these results suggest a multi-dimensional effect of secondary variants towards clinical features, and that their contribution to specific phenotypic domains depends on the extent to which the primary variant sensitizes an individual towards a specific phenotypic trajectory. An important observation from our study is that a large number of disease-associated variants that were deemed to be solely causative for the disorder are in fact accompanied by substantial amount of rare genetic variation. Longitudinal and quantitative phenotyping across multiple developmental domains in all family members, along with whole genome sequencing studies in affected and seemingly asymptomatic individuals with a primary variant, are necessary for a more accurate understanding of these complex disorders. Therefore, it is critical that even after identifying a likely diagnostic disruptive mutation, further analysis of the genetic background must be performed in order to provide appropriate counseling and management.

## Acknowledgements

We are grateful to all of the families in each cohort (16p12.1 deletion, SVIP and SSC) who participated in the study. We thank the SSC principal investigators (A. Beaudet, R. Bernier, J. Constantino, E. Cook, E. Fombonne, D. Geschwind, R. Goin-Kochel, E. Hanson, D. Grice, A. Klin, D. Ledbetter, C. Lord, C. Martin, D. Martin, R. Maxim, J. Miles, O. Ousley, K. Pelphrey, B. Peterson, J. Piggot, C. Saulnier, M. State, W. Stone, J. Sutcliffe, C. Walsh, Z. Warren, E. Wijsman) as well as the Simons VIP Consortium. We appreciate obtaining access to genomic and phenotypic data on SFARI Base. Approved researchers can obtain the SSC and SVIP population datasets described in this study by applying at https://base.sfari.org.

This work was supported by NIH R01-GM121907, Brain and Behavior Foundation (NARSAD#22535), SFARI Pilot Grant (SFARI #399894, SG) and resources from the Huck Institutes of the Life Sciences to S.G. L.P. was supported by Fulbright Commission Uruguay – ANII and the Huck Institutes of the Life Sciences. M.J. was supported by NIH T32-GM102057. C.R., L.C., O.G. and E.A. were supported by the Italian Ministry of Health and ‘5 per mille’ funding. K.M. is a Jacobs Foundation Research Fellow. A.R. is supported by the Swiss National Science Foundation (31003A_160203). Dedicated to the memory of Ethan Francis Schwartz, 1996-1998.

